# Bulk degradation of dendritic cargos requires Rab7-dependent transport in Rab7-positive/LAMP1-negative endosomes to somatic lysosomes

**DOI:** 10.1101/215970

**Authors:** C.C. Yap, L. Digilio, L.P. McMahon, A.D.R. Garcia, B. Winckler

## Abstract

Regulation of protein homeostasis (“proteostasis”) is necessary for maintaining healthy cells. Disturbances in proteostasis lead to aggregates, cellular stress and can result in toxicity. There is thus great interest in when and where proteins are degraded in cells. Neurons are very large as well as very long-lived, creating unusually high needs for effective regulation of protein turnover in time and space. We previously discovered that the dendritic membrane proteins Nsg1 and Nsg2 are short-lived with half-lives of less than two hours. Their short half-lives enabled us to ask whether these proteins are degraded by local degradative pathways in dendrites. We discovered a striking spatial gradient of late endosomes/lysosomes in dendrites, with late endosomes (Rab7-positive/LAMP1-negative/cathepsinB-negative) found in distal portion of dendrites, and degradative lysosomes (LAMP1-positive/cathepsinB-positive) being overwhelmingly found in the soma and in the proximal portion of dendrites. Surprisingly, the majority of dendritic Rab7-positive late endosomes do not contain LAMP1, unlike Rab7-positive late endosomes in fibroblasts. Secondly, Rab7 activity is required to mobilize these distal pre-degradative dendritic late endosomes for transport to the soma and degradation. We conclude that the vast majority of dendritic LAMP1-positive endosomes are not degradative lysosomes and that bulk degradation of dendritic cargos, such as Nsg1, Nsg2, and DNER, requires Rab7-dependent transport in late endosomes to somatic lysosomes.

## Introduction

Protein homeostasis and regulation of the half-lives of proteins are essential to cellular health in all cell types. Since neurons are both post-mitotic and long-lived, maintaining the proteome is of particular importance. Not surprisingly, disturbances in protein homeostasis in neurons has been associated with numerous neurodegenerative disorders, as well as aging (Douglas and Dillin, 2010; Cornejo et al., 2017). In addition to being long-lived, neurons are also extraordinarily large cells whose processes, axons and dendrites, can span hundreds of micrometers or more. How protein turnover is regulated in time and space in axons and dendrites is thus an essential cell biological question (Jin et al., 2017).

Nsg1 (=NEEP21) and Nsg2 (=P19) are neuronal-specific, small (21kd and 19kd) transmembrane proteins found specifically in dendritic endosomes. Nsg1 has been implicated in regulating the trafficking of multiple neuronal receptors, including the axonal cell adhesion molecule L1, the neurotransmitter receptor GluA2, and the Alzheimer disease-associated βAPP (Steiner et al., 2005; Yap et al., 2008; Norstrom et al., 2010). Mistargeting of these receptors after knockdown of Nsg1 resulted in shorter axon growth on L1 substrate (Lasiecka et al., 2014), impaired long term potentiation in hippocampal slices (Alberi et al., 2005), and increased amyloidogenic processing of βAPP (Norstrom et al., 2010). While studying the trafficking of Nsg1 and Nsg2 to dendritic endosomes, we discovered that Nsg1 and Nsg2 are short-lived in neurons (Yap et al., 2017). This was very surprising since the receptors regulated by Nsg1 are involved in long-lasting processes, such as axon guidance and synaptic function. The rapid degradative flux of the Nsg proteins allows us to ask where dendritic membrane proteins are degraded and how their degradation is regulated since changes in protein levels can easily be detected in interference experiments.

In fibroblasts, endosomal populations and their maturation are well studied, and several markers are routinely used to distinguish early endosomes (EE) (EEA1/Rab5) from late endosomes (LE)/lysosomes (lys) (Rab7/LAMP1). Rab7 co-localizes extensively with LAMP1 or LAMP2 in fibroblasts (>75%) (Humphries et al., 2011; Hubert et al., 2016), and both Rab7-positive/LAMP1-positive compartments as well as Rab7-negative/LAMP1-positive compartments are terminal compartments for degradative cargos (Humphries et al., 2011). In fact, the overlap of Rab7 with LAMP1 is so extensive in fibroblasts that late endosome and lysosomes are often combined operationally into a single “LE/lys” category. Absence of M6PR is sometimes used as the one criterion that distinguishes LEs from lysosomes. Using many of the same markers as well as functional assays routinely used in fibroblasts, we began a comprehensive categorization of endosomes in neurons. We discovered a striking spatial gradient of endosomes in different stages of maturation along dendrites: early and late endosomes are abundant all along dendrites whereas the vast majority of degradative, acidified lysosomes are excluded from dendrites ex vivo and in vivo. Secondly, we discovered that dendrites contain distinct Rab7-positive compartments which we term “early” LE and “late” LE, depending on whether LAMP1 is present or not. Unlike in fibroblasts, Rab7-compartments in dendrites frequently lack LAMP1, are not acidified, and do not contain cathepsinB. Thirdly, we show that long-range Rab7-dependent, dendritic transport back to the soma (subsequently referred to as “retrograde”) is required for degradation and that bulk degradation does not occur locally in dendrites. We propose that turnover of dendritic membrane proteins in general occurs overwhelmingly in somatic lysosomes and that LAMP1-negative “early” late endosomes are the primary retrograde carrier for dendritic degradative cargos. These observations raise important questions about how overall protein homeostasis is regulated in distal portions of dendrites which largely lack lysosomes.

## Results

### Nsg family proteins and DNER have short half-lives in neurons whereas proteins resident in late endosomes/lysosomes are longer lived

We recently discovered that the endosomal somatodendritic transmembrane proteins Nsg1 and Nsg2 have short half-lives of about 90 minutes in cultured neurons (Yap et al., 2017). Since Nsg1 and Nsg2 showed very similar half-lives, trafficking, and localization to the early-to-degradative arm of the dendritic endosomal system, we refer to them collectively as “Nsg proteins” in this work. We first investigated if proteins that have been linked functionally to Nsg1 are similarly short-lived. Protein levels of L1, AMPAR and syntaxin13 were determined in cultured rat hippocampal neurons at DIV8/9 treated with cycloheximide (CHX) for up to four hours. Remaining protein levels ranged from ~70 to 80% after four hours of CHX for L1, AMPAR (GluA2), and the early/recycling endosome-resident SNARE protein syntaxin 13 (Figure 1A). Since Nsg proteins are found in late endosomes, we then determined if other proteins found in the late endosome/lysosome were short-lived. 90% or more of the initial protein levels remained for the late endosomal SNARE syntaxin 7, and the lysosomal proteases cathepsin (cat) B and cat D (Figure 1B). The levels of the lysosome-associated membrane protein LAMP2 remained at ~80% after four hours in CHX (Figure 1B). In striking contrast, levels of Nsg1 and Nsg2 were reduced to <20% by four hours (Figure 1C; Yap et al., 2017).

**Figure 1:**
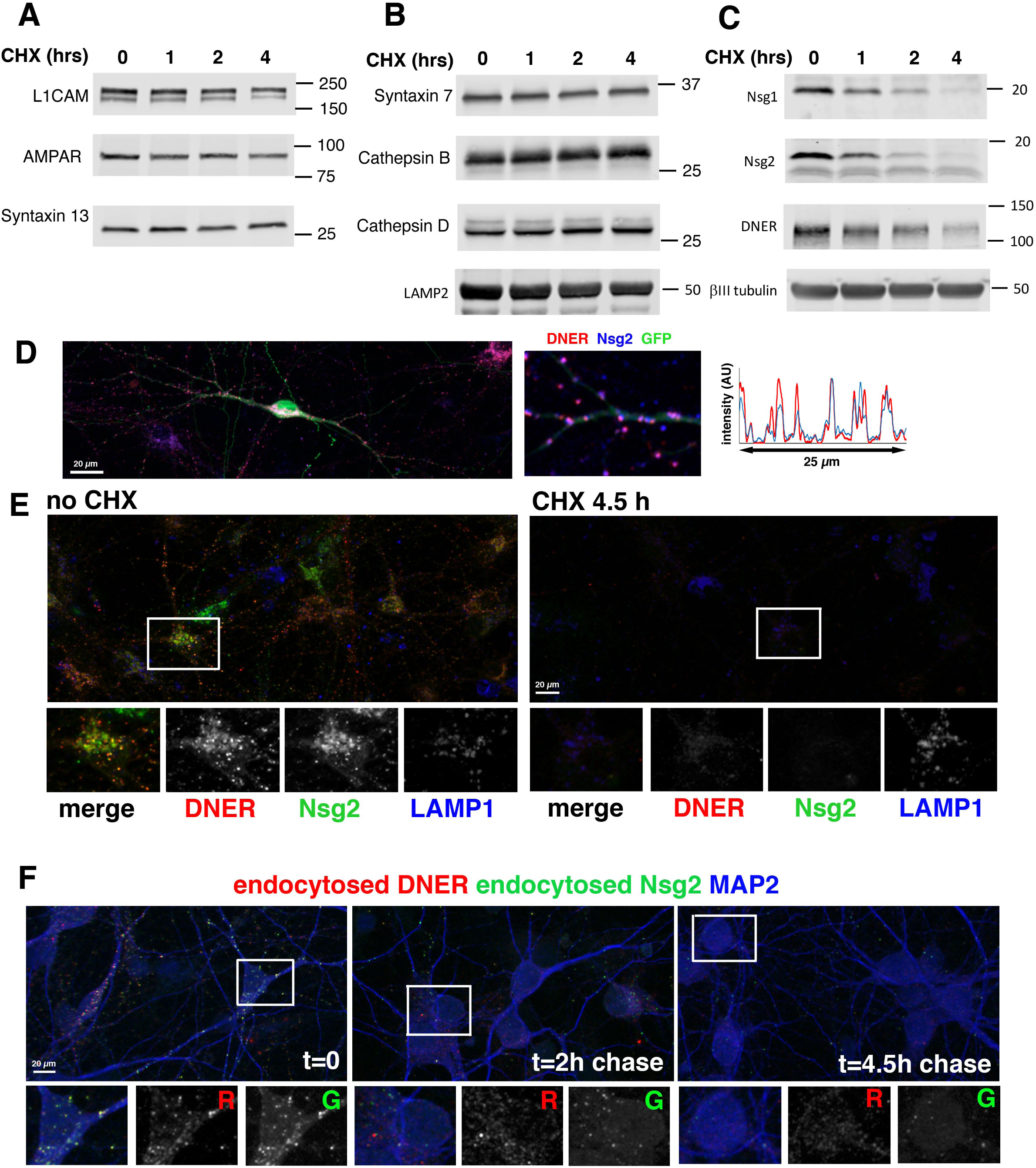
Nsg family proteins and DNER have short half-lives in neurons. (A-C) Protein levels from neuronal cultures were determined after 0, 1, 2, and 4 hours of cycloheximide (CHX) treatment. Membrane proteins associated with the plasma membrane (L1CAM and AMPAR) and recycling endosomes (stx13) were probed in (A), proteins localized to late endosomes and lysosomes in (B), and dendritic membrane proteins with short half lives in (C). Tubulin is shown as an example of a cytosolic protein and used as a loading control. (D) Endogenous DNER receptor largely co-localizes with Nsg2 which localizes to EEs and LEs in dendrites. A line scan is shown to demonstrate the extent of peak co-incidence of DNER and Nsg2. (E) Neuronal cultures were incubated without CHX or with CHX for 4.5 hours and then immunostained against endogenous Nsg2 (green), DNER (red), LAMP1 (blue) and MAP2 (not shown). DNER and Nsg2 levels are greatly decreased after 4.5 h of CHX treatment whereas LAMP1 levels are largely unchanged. The boxed region is shown below as merged and single channels. Size bar = 20 *µ*m. (F) Degradation of DNER and Nsg2 takes place after endocytosis from the surface. Neuronal cultures were incubated live simultaneously with antibodies against extracellular epitopes of Nsg2 and DNER and cultures fixes immediately (t=0), or after 2h or 4.5h chase at 37°C. The endocytosed pool becomes fainter by 2 hours of chase and is greatly diminished after 4.5 hours of chase. This agrees with the time courses shown in (C) and (E). R= red channel. G = green channel. One example is boxed and shown below as merged and single channels. Size bar = 20 *µ*m.

In order to identify additional short-lived membrane proteins, we mined the literature and found that proteomic studies previously revealed that both Nsg1 and the dendritic membrane receptor DNER (delta-notch EGF repeat containing receptor) accumulated highly in cathepsin B/L double knock out mice, suggestive of high turnover rates (Stahl et al., 2007). We found that DNER indeed had a short half-life of about 2.5 hours in hippocampal neurons (Figure 1C). Endogenous DNER was localized to puncta in the soma and dendrites, consistent with previous work (Eiraku et al., 2002; 2005). Nsg2 localized primarily to early and late endosomes in dendrites (Yap et al., 2017) and DNER greatly co-localized to the same compartments (Figure 1D).

In order to more directly compare the turnover rates of LAMPs, Nsg, and DNER specifically in neurons, we used immunofluorescence microscopy and identified neurons by MAP2 counterstain. Large differences of protein stability in the presence of CHX between endogenous Nsg2, DNER, and LAMP1 were also observed with this assay (Figure 1E): LAMP1 staining remained easily detectable after 4.5h of CHX whereas both Nsg2 and DNER levels were greatly reduced. We also carried out endocytosis assays using antibodies against the extracellular domain of endogenous DNER. The levels of endocytosed DNER were greatly decreased after a 4.5h chase (Figure 1F), similarly to simultaneously endocytosed Nsg2. DNER is thus also an itinerant rather than resident endosomal protein which travels to lysosomes for rapid degradation after endocytosis, similarly to Nsg proteins (Yap et al., 2017).

### Early and late endosomes are distributed all along the lengths of dendrites

Since some membrane proteins which localized to dendrites had very short half-lives (such as Nsg proteins and DNER), we wondered if degradation was taking place locally in dendritic lysosomes. We first determined the spatial distribution of EEs and LEs in dendrites (Figure 2A-C) using endogenously expressed EEA1 (EE), Rab7 (LE), and Nsg2 (EE+LE) as markers. MAP2 counterstaining was used to specifically demarcate dendrites (Figure 2A-C). All markers stained abundant puncta in the soma (inset box 1), but also extended a significant distance into dendrites (inset region 2) (Figure 2B), often being detectable more than 100 *µ*m past the soma to near the distal tips (inset region 3) (Figure 2C). The staining intensity of many endosomes tended to decline somewhat with farther distance from the soma (probably mostly a reflection of their smaller size), but many bright compartments were consistently observed in the distal dendrite (Figure 2 B-C). EEA1 and Rab7 showed partial co-localization: ~30% of Rab7 co-localized with EEA1 and ~20% of EEA1 co-localized with Rab7, consistent with the notion that Rab7 is first recruited onto early endosomes and then triggers their conversion to late endosomes (Rink et al., 2005). The dually positive EEA1/Rab7 compartments therefore likely correspond to transitioning early/late endosomes, in agreement with our previous observations of stationary early endosome converting to late endosomes locally in dendrites (Lasiecka et al., 2014).

**Figure 2:**
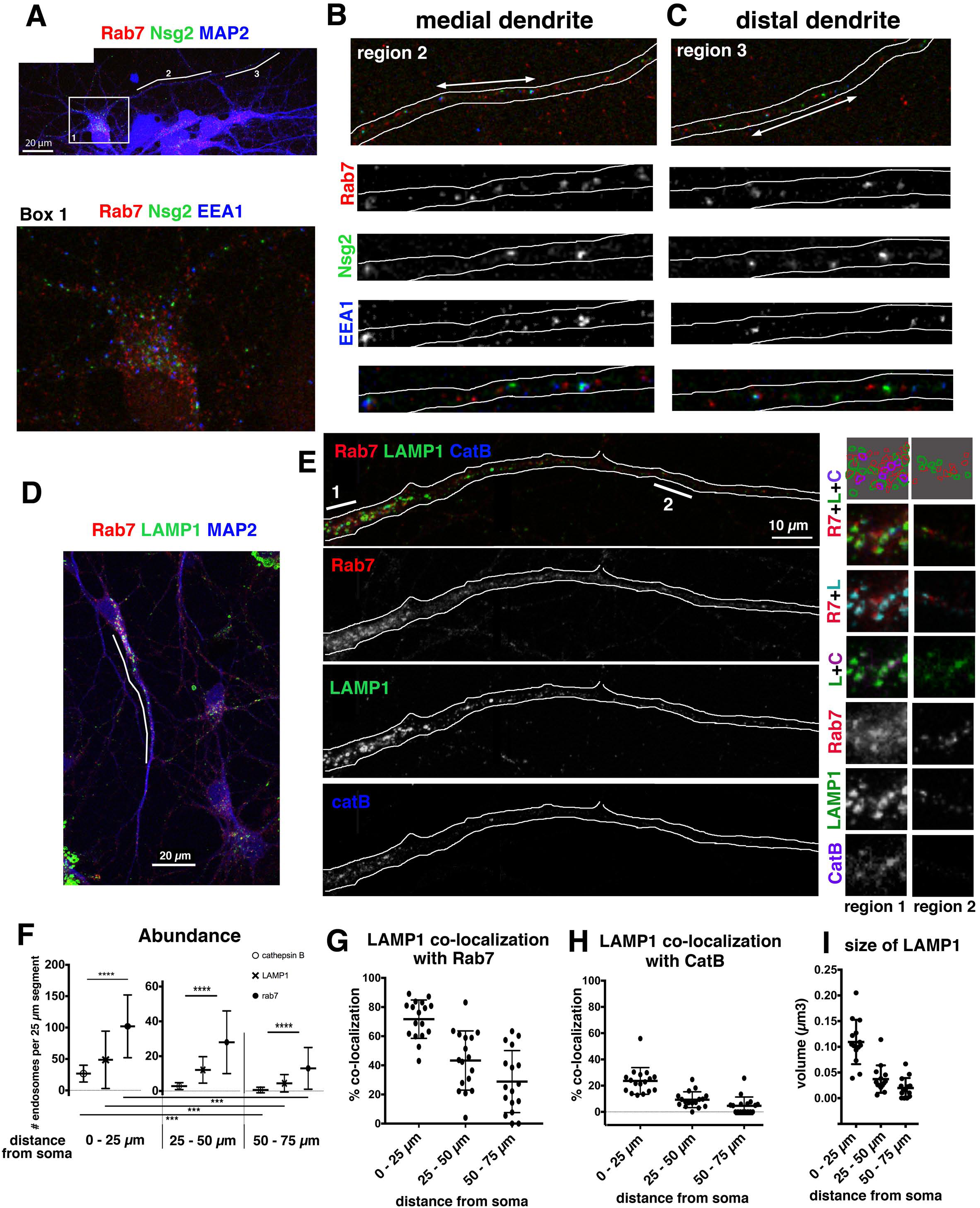
LAMP1-positive endosomes in dendrites rarely contain cathepsin B (CatB) past the proximal region. (A-E) Early (EEA1) and late endosomes (Rab7) (A-C) extend far into dendrites whereas LAMP1- and CatB-positive compartments (D,E) are less abundant past 50 *µ*m from the soma. (A-C) DIV9 hippocampal neurons were stained against MAP2 (to mark dendrites), EEA1, Nsg2, and Rab7 (all endogenous proteins). A zoomed in panel of the soma region is shown in Box1, regions 2 and 3 are shown as merged channels in (B) and (C), respectively. The position of regions 2 and 3 are indicated by a white line in (A). A close-up of the segments marked by double-headed arrows in (B) and (C) are shown as individual channels below. Positively staining endosomes for all three markers can be observed all along the dendrite. (D,E) DIV7 hippocampal neurons were stained against MAP2 (to mark dendrites), Rab7, LAMP1 and CatB (all endogenous proteins). An overview of several neurons is shown in (D). The first hundred micrometers of the largest dendrite (marked by white line) is shown as merged channels and single channels in (E). Rab7 extends far into the dendrite, whereas CatB is largely restricted to the first 25 *µ*m past the soma. LAMP1-positive compartments extend past the first 25 *µ*m, but are less frequent and smaller more distally. Close-ups of the regions marked “1” (proximal) and “2” (medial) are shown in the right hand panels. Single channels and merged channels are shown as indicated (L=LAMP1, R7=Rab7, C=CatB). A schematic representation of the compartment subtypes is shown in the top panels. Purple compartments = degradative lysosomes = LAMP1 + CatB. Green plus red compartments = late LE = LAMP1 + Rab7. Red alone compartments = early LE = Rab7, but no LAMP1. Green alone compartments = uncharacterized “LAMP only” compartments, not LE, not lysosome. See Suppl. Fig. 1 for Rab7-GFP in combination with endogenous EEA1 and LAMP1. (F-I) Quantification of the distribution of endogenous Rab7, LAMP1, and CatB from confocal z-stacks. N for panels F-I: Z-stacks from 14-17 DIV7 neurons were reconstructed in 3D and analyzed using Imaris. Many endosomes (50 to >200 endosomes per cell) were averaged for each cell. (F) All markers show the highest abundance in the soma-near 25 *µ*m (proximal) compared to the segments 25-50 *µ*m (juxtaproximal) or 50-75 *µ*m (medial) past the soma (Mann Whitney test). In all segments, there are significantly more Rab7 endosomes compared to LAMP1 and CatB (Kruskal Wallis test). CatB-positive endosomes are rarely observed past the first 50 *µ*m. (G,H) Co-localization of LAMP1 with Rab7 (G) and CatB (H) was quantified in the segments located 0-25 *µ*m (proximal), 25-50 *µ*m (juxtaproximal), and 50-75 *µ*m (medial) past the soma. Co-localization of LAMP1 with either Rab7 or CatB was highest near the soma and was near zero for CatB more distally in most cells. (I) Quantification of the apparent volume of LAMP1-positive compartments reveals that soma-near compartments are larger than more distally positioned compartments. See Figure 9 for summary.

### Distinct spatial distribution of late endosome/lysosome markers along dendrites

Next, we determined the localization of late endosomes and lysosomes in dendrites. We used multiple markers (Rab7, LAMP1, CatB) (Figure 2, Suppl. Fig.1) and functional assays (Lysotracker, MagicRed cathepsin substrates) (Figure 3, Suppl. Figs. 2,3) to identify lysosomes. We define lysosomes as acidified LAMP1-positive compartments containing active cathepsins and employ the term “degradative lysosome” to designate these compartments. We stained against MAP2 (to identify dendrites) together with endogenous Rab7 (LE), CatB (lys), and LAMP1 (LE+Lys) (Figure 2D,E). The abundance and intensity profiles changed profoundly with distance from the soma and was strikingly different for CatB compared to LAMP1 and Rab7. In order to quantify the marker distribution along dendrites (Figure 2F), we subdivided the major dendrite into three regions: 0 - 25 *µ*m from soma boundary = proximal dendrite, 25 - 50 *µ*m from soma boundary = juxtaproximal dendrite, 50 - 75 *µ*m from soma boundary = medial dendrite. LAMP1-containing compartments were present into and past the juxtaproximal dendrite (>50 *µ*m from soma), but were sparser medially compared to proximal dendritic regions. In contrast, CatB staining was almost completely undetectable past the proximal region (Figure 2E). Many of the major dendrites extended distally past 75 *µ*m and were often much longer than 100 *µ*m. LAMP1 and CatB puncta were essentially absent in these distal regions and thus not included in the graph (see Figure 9 for nomenclature). We note that there was a striking difference between dendrites of different diameters. The major dendrite usually contained many CatB-positive compartments in the proximal segment whereas minor dendrites usually showed essentially no LAMP1 or CatB staining even proximally. EEA1, Nsg proteins and Rab7 were all present in minor as well as major dendrites.

**Figure 3:**
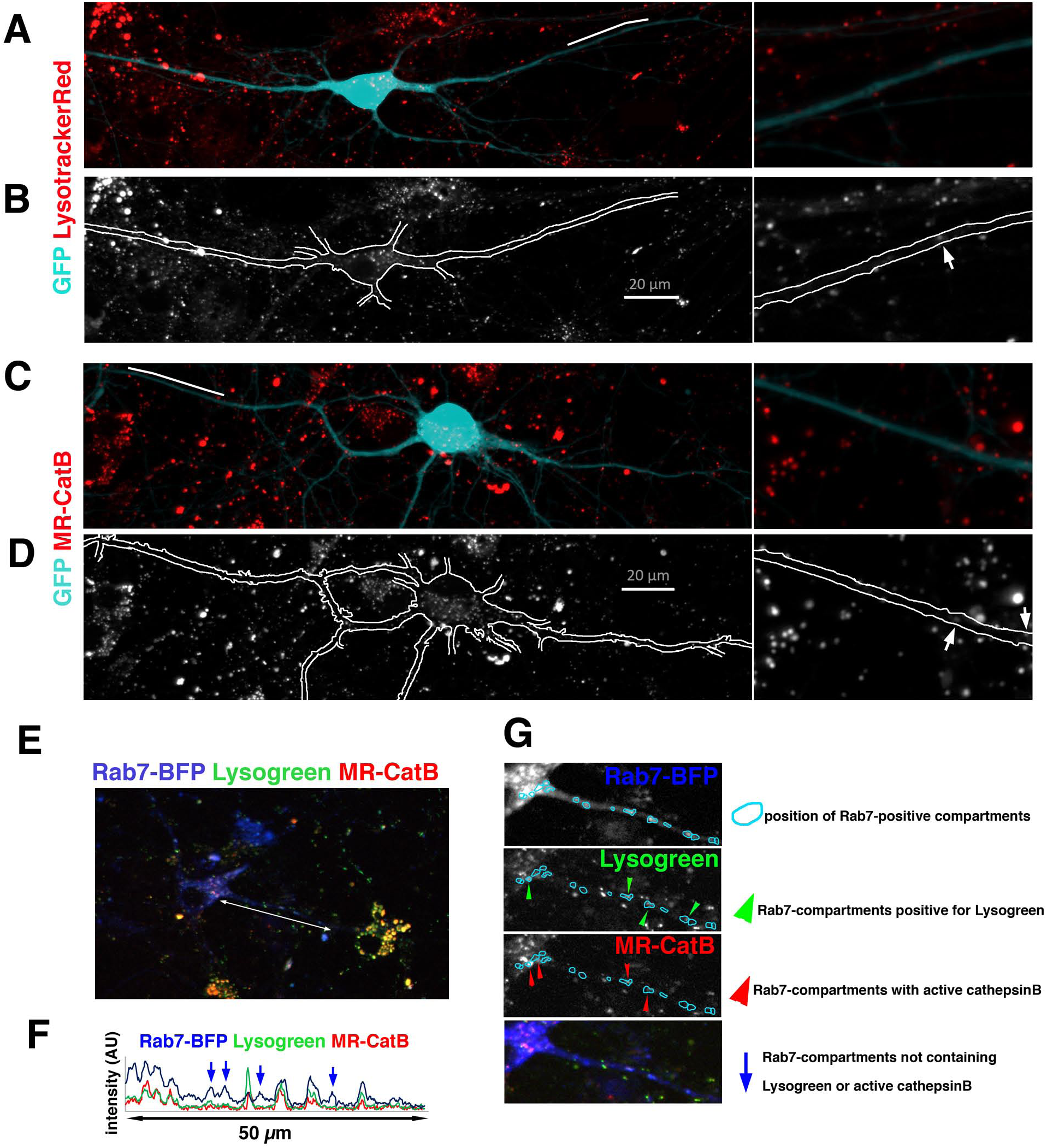
Medial/distal portions of dendrites overwhelmingly lack acidified, cathepsin-containing lysosomes in dissociated neuronal cultures. (A-D) DIV9 neurons were transfected with GFP to show cell morphology and to distinguish dendrite staining from staining contributed by non-transfected cells in the culture. Cultures were imaged live with LysotrackerRed (A,B) or MagicRed-cathepsin B substrate (MR-CatB) (C,D). (A) and (C) show a merged image of GFP (aqua) with the red vital tracer (red). (B) and (D) show single channel image of the red channel only. LysotrackerRed (B) and MR-CatB (D) are most abundant in the soma. The region indicated by a white line in (A) and (C) are shown as close-ups in the right hand panels. Arrows point at the occasional tracer-containing compartments present in more distal parts of dendrites. See Suppl. Figure 2 for co-imaging of endosomes with LysotrackerRed and MR-CatB. See Suppl. Figure 3 for imaging of active cathepsins K and L (MR-CatK and MR-CatL). (E-G) Acidification and presence of active cathepsin B was simultaneously imaged by incubating DIV9 neuronal cultures with Lysogreen (green) and MR-CatB (red) after transfection with Rab7-BFP (blue) to mark late endosomes in live cells. (F) is a line scan along the white line shown in (E). Blue arrows in the line scan indicate non-acidified, CatB-inactive, Rab7-positive endosomes. (G) The soma and proximal dendritic regions of the cell in (E) are shown as single as well as merged channels. Rab7-BFP positive late endosomes are encircled (aqua) and overlaid over each channel to allow comparison of staining. Rab7-BFP positive endosomes that also contain Lysogreen or MR-CatB are indicated by green and red arrowheads, respectively, as indicated in the legend.

### Early late endosomes (Rab7-positive/LAMP1-negative) are abundant in medial and distal dendrites

Quantification of marker abundance in different dendrite regions (Figure 2F) revealed that Rab7-positive compartments are much more abundant in dendrites than LAMP1-positive compartments, and that Rab7-positive compartments frequently did not contain LAMP1. We thus wanted to determine if the Rab7-positive/LAMP1-negative compartments in dendrites were transitioning early endosomes (i.e. EEA1-positive) or constituted a Rab7-positive/EEA1-negative/LAMP1-negative compartment. Due to antibody compatibility constraints, we had to overexpress Rab7 for this marker combination. In the soma, Rab7-GFP greatly co-localized with LAMP1. In contrast, Rab7-GFP compartments in the distal half of dendrites frequently contained neither EEA1 nor LAMP1 (Suppl. Figure 1A,B). We categorize these Rab7-positive/LAMP1-negative endosomes as “early” LEs. Quantification revealed that the co-localization of LAMP1 with Rab7 also decreased with increasing distance from the soma (Figure 2G), such that a large proportion of LAMP1-compartments in the medial/distal dendrites lacked Rab7. The exact identity of these compartments is not clear.

### LAMP1-positive endosomes in dendrites rarely contain cathepsinB

In order to define compartments as degradative lysosomes we determined the extent of LAMP1-positive compartments that also contained lysosomal proteases (such as cathepsins). We stained against endogenous CatB, but note that the antibody does not distinguish inactive preprocessed forms of CatB from cleaved active CatB. We find that LAMP1-positive compartments in the soma and proximal portion of major dendrites co-localized substantially (but partially) with CatB (Figure 2H), making them degradative lysosomes. In contrast, LAMP1-positive compartments further than 25 *µ*m from the soma boundary rarely contained CatB (Figure 2H). These dendritic LAMP1-containing compartments thus might be non-degradative “late” LEs rather than lysosomes with degradative capacity. Rarely, we observed an occasional CatB-positive compartment more distally (>75 *µ*m) in a major dendrite, suggesting limited, local degradative capacity in dendrites past the proximal region. We note that CatB-positive compartments were virtually never observed in major dendrites past 100 *µ*m away from the soma or in minor dendrites at any distance. In addition, the volume of the LAMP1-stained compartments decreased strikingly past the first 25 *µ*m from the soma (Figure 2I). The large LAMP1-compartments appeared “doughnut-shaped” by high resolution confocal microscopy, also typically seen for lysosomes in non-neuronal cells (Suppl. Fig. 1C, arrowheads). LAMP1-compartments more distally in dendrites were small and a lumen could not be resolved (Suppl. Fig. 1C, arrows). We thus discovered a striking spatial gradient of endosomal compartments in dendrites, with early (EEA1-positive) and late endosomes (Rab7-positive/CatB-negative) extending far into the distal portion of dendrites and compartments containing lysosomal proteases (LAMP1-positive/CatB-positive) being most concentrated in the soma and the proximal portion of dendrites.

### Dendrites overwhelmingly lack acidified lysosomes containing active cathepsins

Since lysosomes contain many hydrolases other than CatB, we also used two operational assays to define degradative lysosomes. Lysotracker was used to stain acidified compartments and MagicRed-cathepsin substrates (MR-Cat) to stain compartments that contain active cathepsins. Since both lysotracker and MR-Cat are poorly fixable, we carried out imaging in live cells using GFP to visualize cell shape. Both lysotracker (Figure 3A,B) and MR-CatB (Figure 3C,D) were present abundantly on brightly staining compartments in the soma and proximal portion of the major dendrites, but only an occasional small, faint compartment was visible further out in the dendrites (arrows in Figure 3B,D). In addition, we visualized dendritic endosomes by transfecting Nsg1-Em and imaged LysotrackerRed or MR-CatB (Suppl. Figure 2). Many of the Nsg1-Em containing endosomes near the soma were brightly stained with Lysotracker or MR-CatB, but past the proximal segment the tracer staining became very faint in Nsg1-Em endosomes and less abundant overall. We note that Lysotracker and MR-CatB staining in non-neuronal cells present in our cultures was much stronger than in neurons and was the most easily visible signal in the field.

In order to distinguish degradative lysosomes (defined as simultaneously acidified and degradative) from “early” LEs operationally, we carried out triple live imaging with neurons transfected with Rab7-BFP, and incubated with Lysogreen, and MR-CatB (Figure 3E-G). As expected, MR-positive compartments were acidified (Lysogreen-positive) (see arrowheads in Figure 3G), allowing their identification as degradative lysosomes. Many Rab7-BFP containing compartments, in contrast, did not stain with either Lysogreen or MR-CatB (blue arrows in Figure 3F). We categorize these Rab7-endosomes operationally as “early” LEs (i.e. non-acidified, non-degradative).

Since lysosomes contain many catabolic enzymes including other cathepsins, we additionally stained live neurons with MR-CatL and MR-CatK. Their staining intensity along dendrites also showed a steep decline with increased distance from the soma (Suppl. Figure 3), demonstrating that three different active cathepsins (B, K, and L) all showed a similar steep spatial gradient along dendrites. These functional assays thus led to the same conclusion as the antibody staining, namely that acidified, proteolytically active compartments (i.e. degradative lysosomes) were found mostly in the soma and proximal portion of the major dendrite, were sparsely present in the juxtaproximal dendrite and virtually absent from medial and distal dendrites.

### Rab7 function is required for transport of Nsg proteins

Interference with Rab7 function (either by short hairpin (sh)-based knockdown of Rab7 or by expression of the dominant-negative Rab7-T22N, “Rab7-DN”) led to massive accumulation of Nsg proteins in dendritic endosomes (Yap et al., 2017), suggesting that degradation was impaired in the absence of functional Rab7. Since we observed very limited degradative capacity localized to dendrites past the proximal dendrite but abundant Rab7-positive endosomes (Figure 2 and 3), we hypothesized that Rab7 was required for transport of pre-degradative dendritic endosomes to the soma. GFP (Figure 4A) or WT Rab7-GFP (Figure 4B) were used as controls, and Rab7-DN was used to interfere with Rab7 function (Figure 4 C-E). We first determined if Nsg proteins accumulated in EEs (EEA1; Figure 4C) or dendritic LEs (LAMP1; Figure 4D) when Rab7-DN was expressed. As we found previously (Lasiecka et al., 2014; Yap et al., 2017), Nsg proteins showed partial (~20-30%) co-localization with EEs (Figure 4A) and with Rab7-GFP (Figure 4B2, green arrowheads) including dually positive endosomes (EEA1/Rab7-GFP; aqua arrowheads), representing transitioning EEs. When Rab7-DN was expressed (Figure 4C), Nsg accumulated 2-3 fold higher than in controls throughout dendrites, including in the distal portion of long dendrites (Figure 4C3). A subset of these accumulated compartments contained EEA1 (blue arrows; Figure 4C1, C2). EEA1-positive endosomes did not change either intensity or abundance in Rab7-DN expressing neurons.

**Figure 4:**
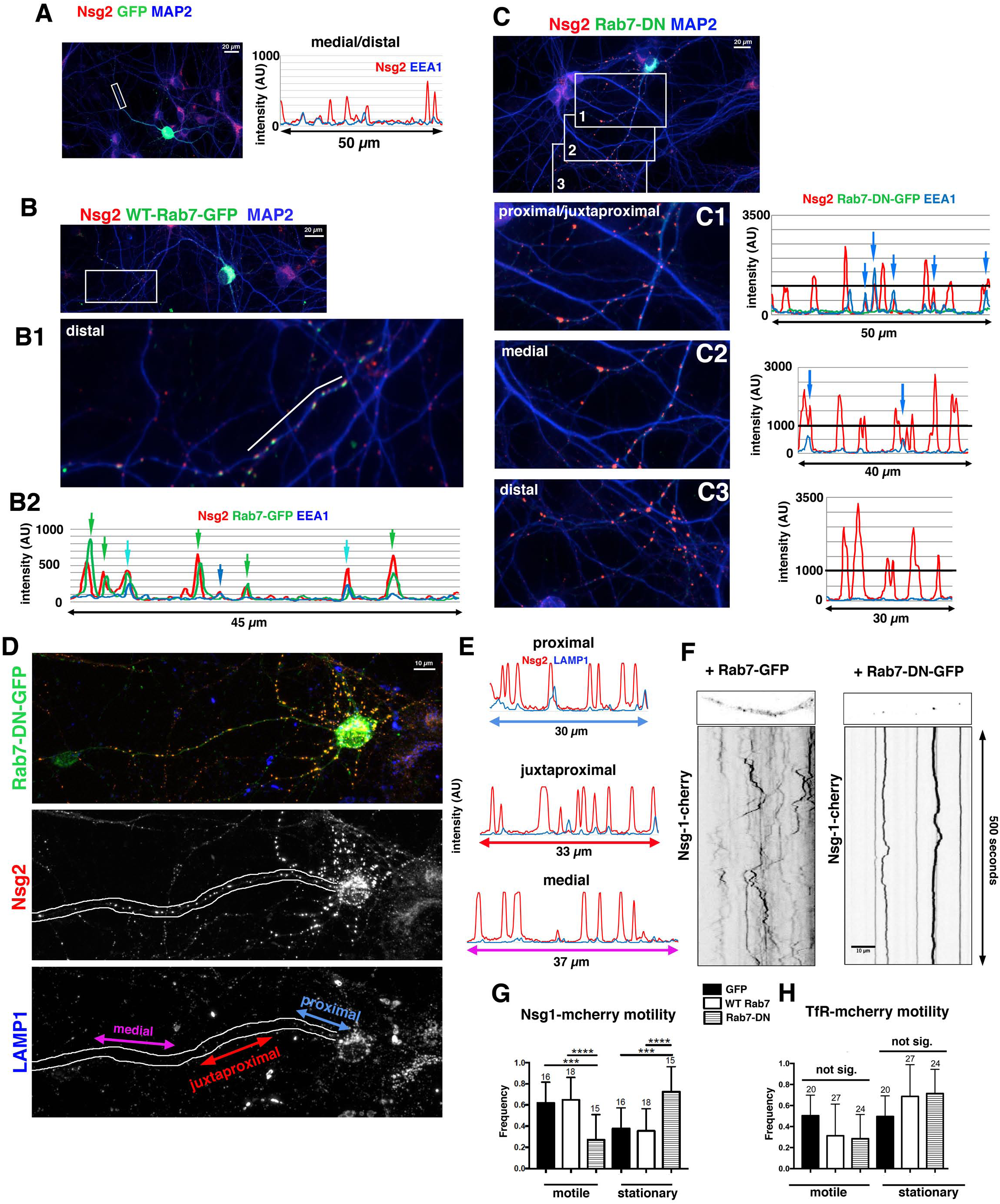
Interference with Rab7 leads to accumulation of the short-lived dendritic Nsg and DNER proteins along dendrites and cessation of transport. (A-E) Interference with Rab7 leads to accumulation of endogenous Nsg2 (red) along dendrites, including medial/distal regions (>75 *µ*m from soma). (A-B) Expression of WT Rab7-GFP (B) does not change Nsg2 (red) intensity along dendrites compared to GFP expression (A). Nsg2 (red) is found partially in EEA1-positive (blue) compartments (see coinciding peaks in the line scan). When Rab7-GFP is expressed (B), Nsg2 along dendrites is found with EEA1 (blue arrow in B2), with Rab7-GFP (green arrows in B2), or in compartments dually positive for EEA1 and Rab7-GFP (aqua arrows in B2). The intensity of Nsg2 peaks is in the same range as for GFP expression (A). The location of the line scan in B2 is indicated by a white line in B1. B1 corresponds to the boxed region in B. MAP2 (blue) is counterstained to identify dendrites. (C-D) Expression of Rab7-DN-GFP leads to greatly increased Nsg2 levels along dendrites. (C) Neurons expressing Rab7-DN-GFP show accumulation of endogenous Nsg2 (red) along dendrites (marked with MAP2 in blue). The proximal/juxtaproximal (C1), medial (C2), and distal (C3) dendritic regions (boxed in panel C) are shown with corresponding line scans. Exposures were identical to panels A and B for direct comparison of peak heights. The dark line on the line scans in C1-C3 corresponds to 1000 AU intensity for easier comparison to line scans in A and B2. Nsg2 accumulation partially occurs in EEA1-positive early endosomes (blue arrows in line scans). In C3, accumulated Nsg2 is not in EEA1-positive compartments. DNER also accumulates highly along dendrites in neurons expressing Rab7-DN-GFP (shown in Suppl. Figure 4). (D,E) Nsg2 (red) accumulates partially in LAMP1-positive (blue) compartments in proximal/juxtaproximal dendrite segments in cells expressing Rab7-DN-GFP (see linescans in E). The line scans shown in E are marked by double-headed color-coded arrows in D. Unlike Nsg2, LAMP1 levels are not changed in Rab7-DN-GFP expressing cells compared to untransfected neurons in the same field. (F-H) The motility of Nsg1-mcherry is significantly inhibited in cells expressing Rab7-DN-GFP compared to GFP or WT Rab7-GFP. Examples of kymographs are shown in F. Quantification is shown in G. (H) To determine the mobility of a non-degradative cargo, TfR-mcherry was co-expressed with GFP, WT Rab7-GFP, or Rab7-DN-GFP and imaged live. No statistically significant changes in TfR-mcherry mobility was observed. Numbers above the bars are N= # of cells. For each cell, 4-5 dendrites on average were counted and averaged to derive a single number per cell. Total dendrite numbers were thus 52-62 for Nsg1 movies and 110-140 for TfR movies. Statistical analysis was done on a per cell basis. Cells were imaged from at least 3 independent cultures. Statistical test was One-way ANOVA (Nsg1) and Kruskal-Wallis (TfR). p<0.0001.

In contrast to Nsg proteins, the abundance, intensity and localization of LAMP1 was unaltered in neurons expressing Rab7-DN, and LAMP1 compartments tapered in abundance and brightness in dendrites (Figure 4D), similarly to controls. In the proximal dendrite, Nsg accumulated partially with LAMP1. Similarly, in the juxtaproximal dendrite where LAMP1 is still easily detected, accumulated Nsg partially co-localized with faint LAMP1-compartments. In contrast, in medial dendrites and beyond, few if any compartments with accumulated Nsg contained LAMP1 (Figure 4E). These observations suggest that Rab7-DN led to accumulation of Nsg distally in pre-degradative (LAMP1-negative) compartments. We posit that these distal compartments include transitioning EEs as well as “early” LEs.

Nsg proteins in dendrites frequently move together with Rab7-positive LEs, but rarely with Rab5-positive EEs (Yap et al., 2017). Since Rab7-DN no longer binds to its downstream effectors (Guerra and Bucci, 2016), including to adaptors for microtubule motors, we determined by live imaging if expression of Rab7-DN impeded the motility of endosomes containing Nsg1-Em. GFP and WT-Rab7 were expressed as controls. Nsg1-Em-positive carriers were frequently motile in controls, but virtually stopped all movements in Rab7-DN expressing neurons, with only short bi-directional excursions (Figure 4F,G). In order to determine if the motility of other endosomes was also slowed by expression of Rab7-DN, we carried out live imaging of TfR-cherry (TfR = transferrin receptor) which is transported in recycling endosomes. No differences of TfR-cherry motility were found between WT Rab7 and Rab7-DN (Figure 4H), consistent with previous findings in other cell types (Saxena et al., 2005). Interference with Rab7 function thus preferentially interfered with transport of cargos fated for degradation.

### Rab7 function is required for degradation of dendritic proteins

Our data suggest the following model: short-lived dendritic membrane proteins rapidly enter Rab7-positive/LAMP1-negative pre-degradative “early” LEs in dendrites and are transported retrogradely in LEs until they reach the proximal dendrite and soma where they fuse with lysosomes. Failure to transport LEs from dendrites to the soma (as occurs in Rab7-DN) is thus predicted to change their half-lives. We used CHX (as in Figure 1E) to test this notion. Neither expression of GFP (Figure 5A) or WT Rab7-GFP (Figure 5B) (green arrow) changed the levels of Nsg2 in untreated controls or after four hours of CHX treatment compared to non-transfected neurons in the same field (aqua arrowheads): Nsg2 degraded similarly and was barely detectable after four hours of CHX treatment. In contrast, expression of Rab7-DN-GFP led to accumulation of Nsg2 (green arrow in Figure 5C) in untreated controls. Additionally, levels of Nsg2 remained high even after four hours in CHX whereas non-transfected neighboring neurons had mostly lost Nsg2 staining (aqua arrowheads in Figure 5C). DNER behaved similarly and showed accumulation in dendritic endosomes in the presence of Rab7-DN which persisted even after 4.5 hours of CHX (Suppl. Fig. 4). Again, LAMP1 was only occasionally present at low levels at the distal compartments in which Nsg2 accumulated in the presence of Rab7-DN and CHX (Figure 5D,E; compare red to purple arrowheads). These Nsg2-containing LAMP1-positive compartments were likely degradation-incompetent since Nsg2/DNER-accumulating endosomes in the distal dendrites were largely CatB-negative (data not shown). Proximally, LAMP1 was often included in the Nsg2-accumulating compartments. We posit that the distal Nsg2-accumulating compartments are “early” LEs whereas the proximal ones correspond to “late” LEs. In addition, endocytosed Nsg2 fails to degrade in the presence of Rab7-DN (data not shown), consistent with an essential role of Rab7 for the degradation of dendritic cargos.

**Figure 5:**
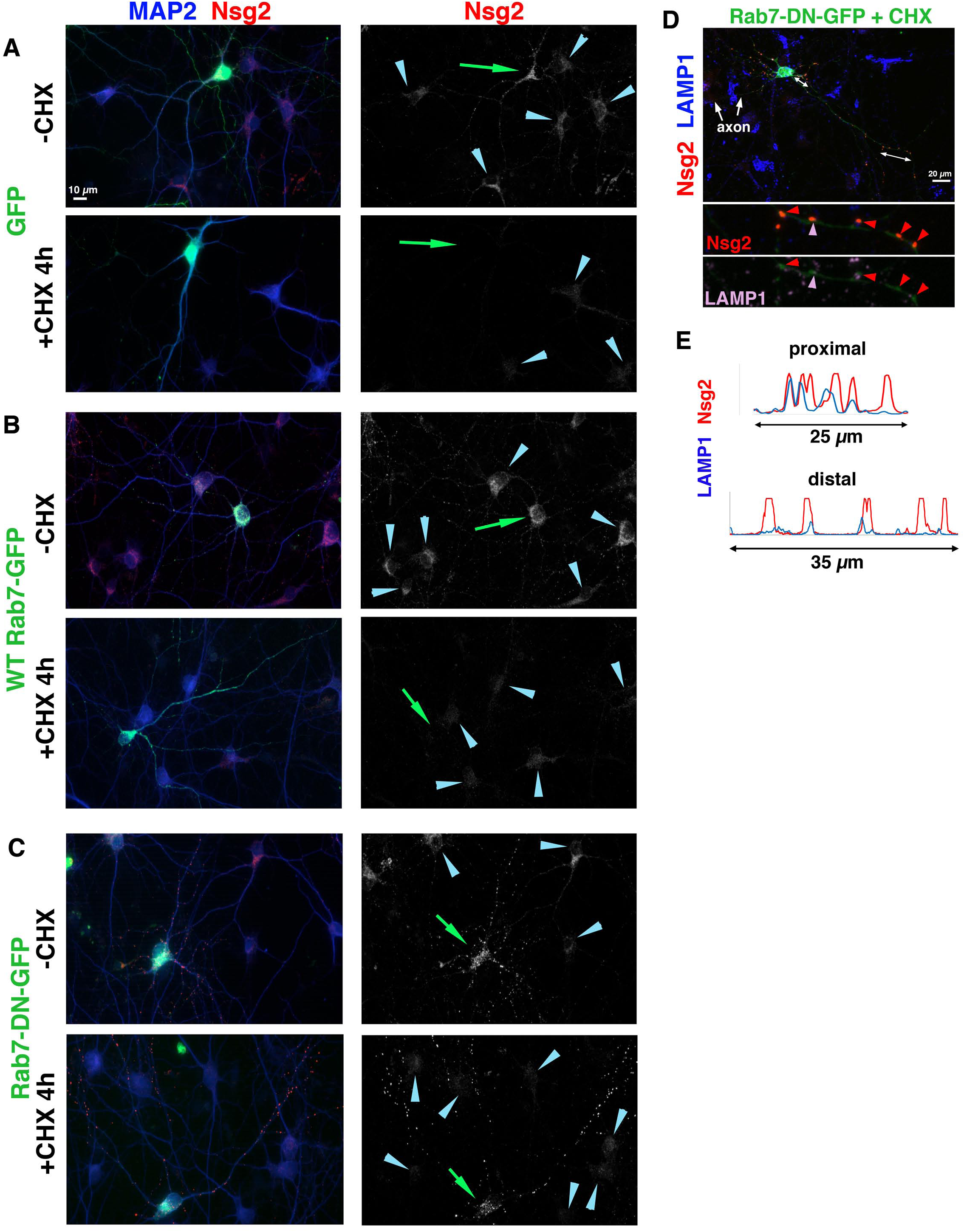
Rab7 function is required for degradation of short-lived dendritic proteins. (A-D) Interference with Rab7 stabilizes Nsg2 in the presence of cycloheximide (CHX). DIV8 cultured neurons were transfected with GFP (A), WT Rab7-GFP (B), or Rab7-DN-GFP (C) and either treated with CHX for four hours (+CHX 4 h) or not treated (-CHX), and then fixed and stained for MAP2 (to mark dendrites) and endogenous Nsg2 (red). Nsg2 single channel images are shown on the right. Nsg2 levels are greatly diminished after 4 h CHX treatment in GFP-(A) and WT Rab7-GFP (B) expressing neurons, but Nsg2 persists at high levels in CHX when Rab7-DN-GFP (C) is expressed. Green arrow points at the transfected cell. Aqua arrows point at untransfected neurons in the same field. The same experiment was carried out for endogenous DNER (Suppl. Fig.4). (D) LAMP1 counterstain of a CHX-treated, Rab7-DN-GFP expressing neuron reveals low co-localization of accumulated Nsg2 (red arrowhead) with LAMP1 (purple arrowhead) in distal dendrites (shown as close up in bottom panel). The region of close-up is indicated by a double-headed arrow in the top panel and shown as a linescan in (E) (distal). A linescan along the proximal dendrite is also shown for comparison. Higher co-localization is observed proximally.

### Inhibition of lysosomal proteases leads to preferential accumulation of Nsg proteins in somatic and not dendritic endosomes

In order to test the effects of inhibiting lysosomal degradation, we used Leupeptin (Leu) to inhibit multiple lysosomal proteases (including CatB, H, L) (Figure 6A). Alternatively, cultures were treated with CHX for 20 hours. Long-lived membrane proteins (GluA2, L1, and endosomal syntaxins) showed moderate changes in protein levels when new synthesis was inhibited with CHX for 20 hours (42-75% of initial level remaining). When lysosomal degradation was inhibited with Leu for 20h, no significant accumulation occurred for these proteins (80-100% remaining). In contrast, Nsg proteins were undetectable after 20h of CHX treatment and their levels increased 1.7- to 2-fold after Leu treatment, demonstrating that their steady-state levels are tightly regulated by new synthesis and lysosomal degradation (Figure 6A). Interestingly, the levels of cathepsins were largely unaffected by CHX treatment, but were increased (about 150% of control) after Leu treatment. A second slower-mobility band could be detected in Leu which likely corresponds to uncleaved pro-forms of cathepsins (Figure 6A). By immunofluorescence, we observed a significant increase in somatic Nsg protein staining after Leu treatment of neurons (Figure 6B), but no increase in somatic Rab7 or EEA1 staining (Figure 6C), consistent with the hypothesis that somatic lysosomes were responsible to a large extent for the degradation of these short-lived proteins.

**Figure 6:**
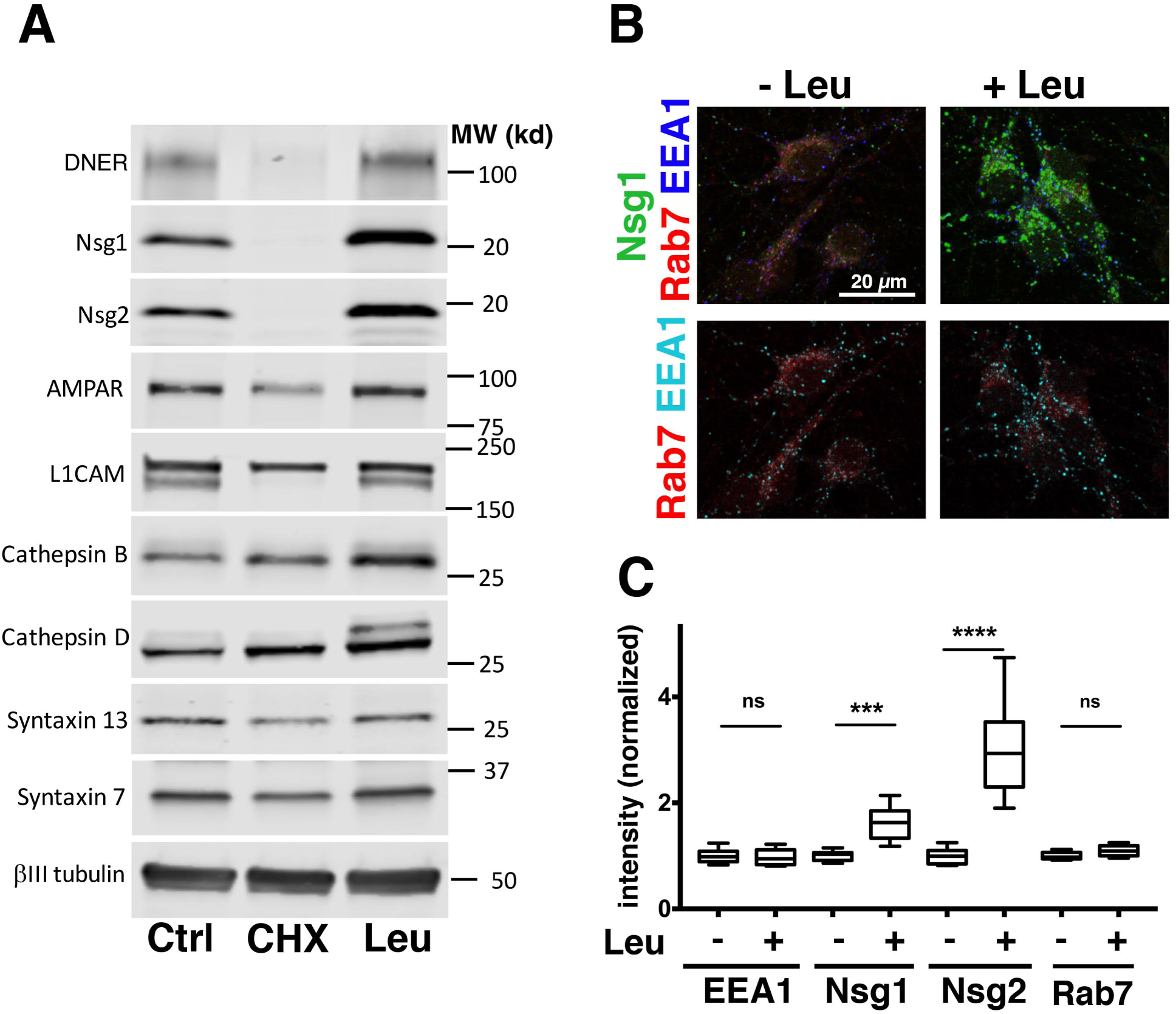
The steady-state levels of Nsg1, Nsg2, and DNER are highly sensitive to inhibitors of protein synthesis or lysosomal proteases. (A) Levels of proteins in neuronal cultures revealed by Western blots after 20 hours of cycloheximide (CHX) or Leupeptin (Leu) treatment. (B,C) Immunostaining of neuronal cultures against endogenous Nsg1 (green), Rab7 (red) and EEA1 (blue) after treatment for 20 hours with and without Leu (20 *µ*m). Quantification of EEA1, Nsg1, Nsg2, and Rab7 in somatic endosomes is shown in (C). 10 fields acquired by confocal microscopy were quantified and values normalized to “no Leu”. The counts for each field included multiple neurons. The data passed a normality test (D’Agostino and Pearson, Prism software). t-test was used to compare with and without Leu for each marker. Only Nsg1 and Nsg2 were significantly accumulated in the soma after Leu treatment. p<0.0001.

We previously found that Nsg2 is endocytosed along dendrites and in the soma, and this endocytosed pool disappeared from both soma and dendrites almost completely when chased for four hours (Yap et al., 2017). We reasoned that if we carried out the Nsg2 endocytosis and subsequent chase assay in the presence of Leu, endocytosed Nsg2 would accumulate in the compartments of the cell where degradation would normally take place in the absence of Leu. This assay thus allowed us to quantify the extent of dendritic vs somatic degradation. If the endocytosed Nsg2 was significantly degraded locally in dendritic lysosomes, we should observe accumulation of non-degraded endocytosed Nsg2 along dendrites after the chase, in addition to the somatic accumulation we already observed for steady-state Nsg2 in the presence of Leu (Figure 6). In controls without Leu, endocytosed Nsg2 was abundantly present in both the soma and dendrites (Figure 7A). After 4.5 hours of chase, no endocytosed Nsg2 remained in either soma or dendrites (Figure 7B). When Leu was included in the experiment, endocytosis of Nsg2 was similar to controls without leupeptin (Figure 7C), but substantial accumulation of endocytosed Nsg2 was observed after 4.5 hours of chase (Figure 7D). In the soma, Leu completely blocked the decrease of Nsg2 intensity (Figure 7E). In contrast, Leu did not block the decrease of Nsg2 intensity that occurs in dendrites (Figure 7F).

**Figure 7:**
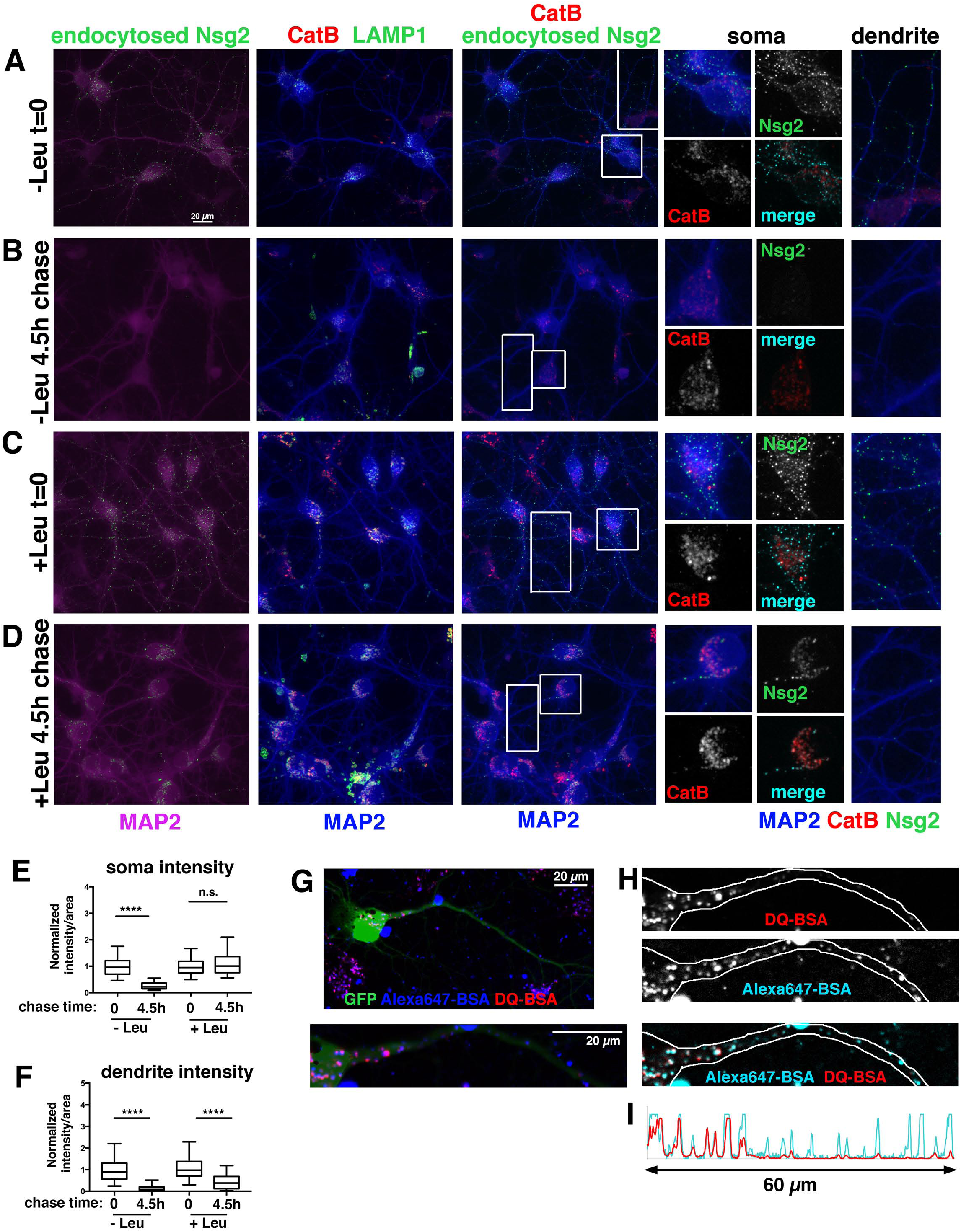
Inhibition of lysosomal proteases leads to preferential accumulation of Nsg proteins in somatic and not dendritic endosomes. (A-D) DIV9 neurons were incubated without (A,B) or with Leu (C,D) and an endocytosis assay against endogenous Nsg2 was carried out. Cells were either fixed directly after antibody loading (t=0; A,C) or endocytosed antibody was chased for 4.5 hours (t=4.5h; B,D). MAP2 is shown in combination with endocytosed Nsg2, endogenous LAMP1 or endogenous CatB as indicated in the panels. MAP2 (blue) with endocytosed Nsg2 (green) and endogenous CatB (red) is shown as close-ups of the soma and dendrite in the small panels on the right. The position of the close-ups are indicated by the white box (soma) and white rectangle (dendrites). A slight increase of the intensity of CatB in neurons treated with Leu was observed. (E,F) The fluorescence intensity of endocytosed Nsg2 in the soma (E) or along dendrites (F) was quantified. Box plots are used to show the 25% and 75% quartile of the data range. The median is indicated. The whiskers show the 5^th^ to 95^th^ percentile of the data range. N= 160-300 cells per condition. Statistical test was Kruskal-Wallis test. P<0.0001. (G-I) Visualization of degradative compartments using internalized DQ-BSA (red). Alexa-647 BSA (blue) was used to visualize all compartments containing BSA. GFP was used to outline the shape of the cell. Single channels are shown in (H) with the corresponding line scan in (I).

### Bulk degradative cargos are preferentially degraded in somatic and proximal dendritic compartments

Our results so far demonstrated that very low cathepsin activity was found past the proximal portion of the major dendrite, but lysosomes do contain many other proteases. We thus used DQ-BSA as a general bulk degradative cargo which turns red upon digestion by any protease, not just cathepsins. DQ-BSA was mixed with Alexa-647 BSA (which is always fluorescent regardless of pH or proteolysis) and fed neurons for two hours (Figure 7G,H). Red fluorescence was brightly visible in the soma and the proximal dendrite and rapidly tapered in abundance and intensity in the juxtaproximal segment. Alexa647-BSA, in contrast, could be brightly detected throughout the dendrite (Figure 7H,I). This experiment thus suggested that bulk degradation overwhelmingly takes place in the soma and proximal dendrite with only low levels of degradative capacity present past the proximal dendrite.

### Lysosomes are rarely found in distal dendrites of mature neurons

We then asked if degradative lysosomes showed a steep spatial gradient in more mature neurons as well. Similar to DIV9 cultures, there were only a few bright LAMP1-positive endosomes in the dendrites of DIV60 neurons, and CatB was rarely detected at all past the proximal region (Figure 8A). Secondly, we stained surface GluA1 to visualize dendrites and spines (Figure 8B). We occasionally detected LAMP1 in a spine close to the soma (similarly to Goo et al., 2017; Padamsey et al., 2017), but CatB staining was not apparent in spines.

**Figure 8:**
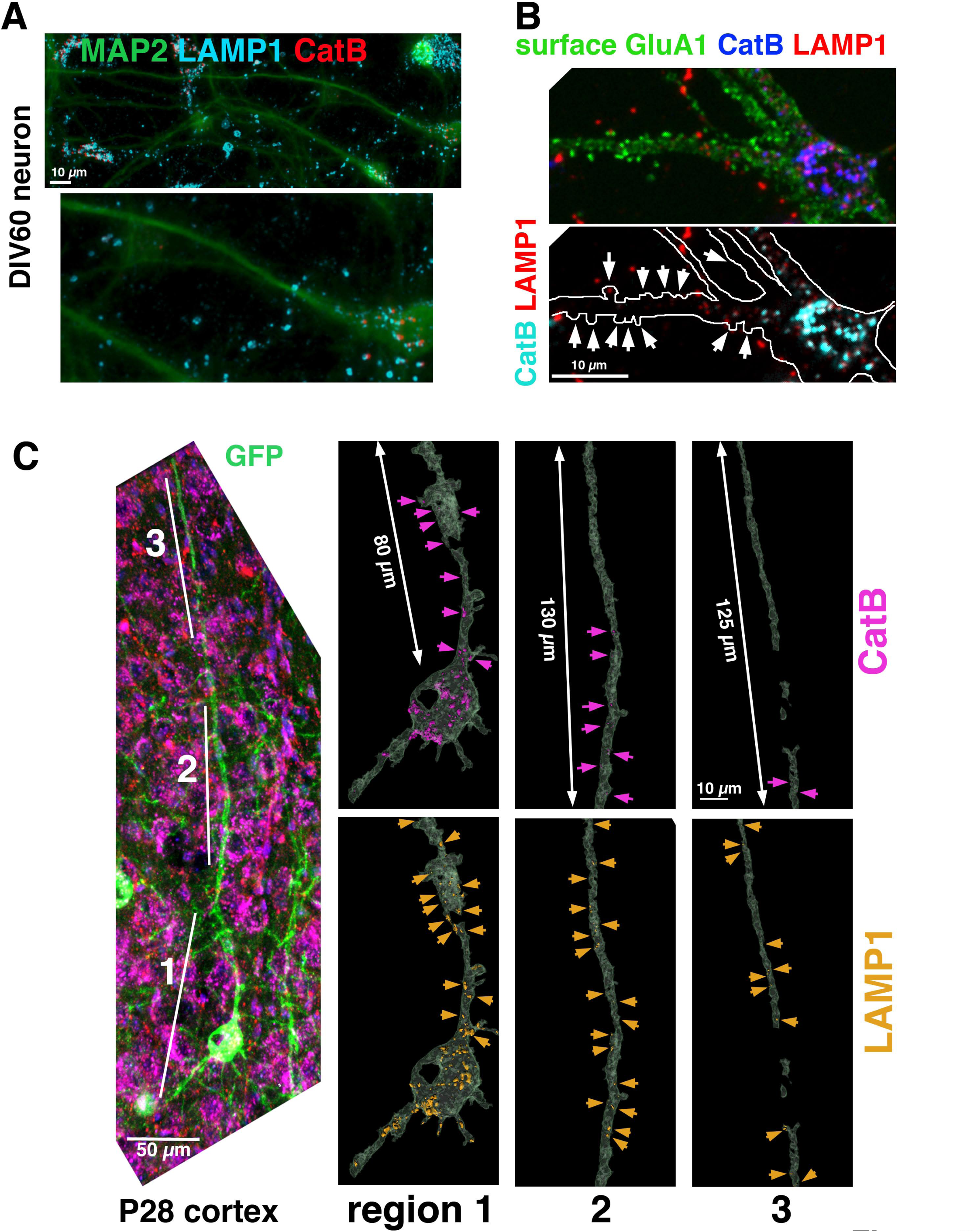
More distal portions of dendrites overwhelmingly lack acidified, cathepsin-containing lysosomes in mature cultured neurons and in mouse cortex. (A,B) Mature dissociated neurons (DIV60) have few cathepsin-positive lysosomes past the proximal dendrite. (A) Neurons cultured for 60 days were stained against endogenous MAP2 (green), LAMP1 (aqua) and CatB (red). (B) Neurons cultured for 60 days were surface stained against endogenous GluA1 (to visualize postsynaptic sites including spines; marked with white arrows) (green), LAMP1 (red), and CatB (blue). We occasionally observed endogenous LAMP1 present in a spine, but did not observe CatB in spines. (C) Coronal cryosections from P28 cortex of mouse brains sparsely expressing GFP (GFP-Thy1 M) were stained against endogenous LAMP1 (red), CatB (pink), and DAPI (blue). The three regions (1-3) marked by white lines in the left panel are shown as reconstructed 3D-rendered single channel images in the right panels. GFP staining was used to mask the dendrite volume (in grey) to allow separation of dendritic signal from signal present in the surrounding cells. LAMP1 (orange) and CatB (pink) positive endosomes contained within the GFP-positive neuron are displayed. Arrows indicate the position of stained compartments. LAMP1-positive compartments (orange arrows) extend well into the dendrite (past 200 *µ*m), but are small past the proximal region. In contrast, CatB-positive compartments (pink arrows) decrease in abundance and size with increasing distance from the soma, and only an occasional CatB-positive compartment of small size can be observed in region 3.

Next, we imaged endogenous LAMP1 and CatB in sparsely GFP-labeled coronal cryosections of P28 mouse cortex using the Thy1-eGFP M line. Neurons whose apical dendrites were largely contained within the cryosection were imaged by confocal microscopy (Figure 8C). LAMP1-containing compartments were abundant in the soma and proximal dendrite, but could also be detected for substantial distances further away towards the more distal end of the dendrite (Figure 8C, orange arrows in left panels). Similarly to cultured neurons, LAMP1-positive compartments past the proximal region were smaller than somatic and proximal LAMP1-compartments. CatB-containing compartments were also abundant in the soma, but then greatly decreased with increasing distance from the soma. CatB was rarely detected in the more distal portions of the dendrite (region 3), and substantial lengths of the medial/distal dendrite contained no compartments containing CatB (Figure 8C, pink arrows regions 2 and 3). We thus find that dendrites of layer2/3 cortical neurons of adult mice contained CatB-containing degradative lysosomes in the first 100 *µ*m segment of the apical dendrite, but CatB-containing degradative lysosomes are rare in the more distal portions of the apical dendrite.

## Discussion

Our recent discovery of the very short half-lives of the somatodendritic endosomal Nsg proteins prompted us to determine where in neurons their degradation was taking place and how it was regulated. We demonstrate here that the overwhelming majority of dendritic LAMP1-positive endosomes are not degradative lysosomes and that bulk degradation of dendritic cargos, such as Nsg1 and Nsg2, requires retrograde Rab7-dependent transport to somatic degradative lysosomes. The same pathway operates for another short-lived dendritic receptor, DNER. DNER is important in cerebellar development, but its exact roles are not well established (Eiraku et al., 2005). In addition, we discovered that medial and distal dendrites have abundant populations of Rab7-positive, LAMP1-negative, lysotracker-negative endosomes which were not previously described in dendrites. We are calling these endosomes “early” LEs and posit that they are maturing LEs transporting to-be-degraded cargos to the soma (Figure 9).

**Figure 9:**
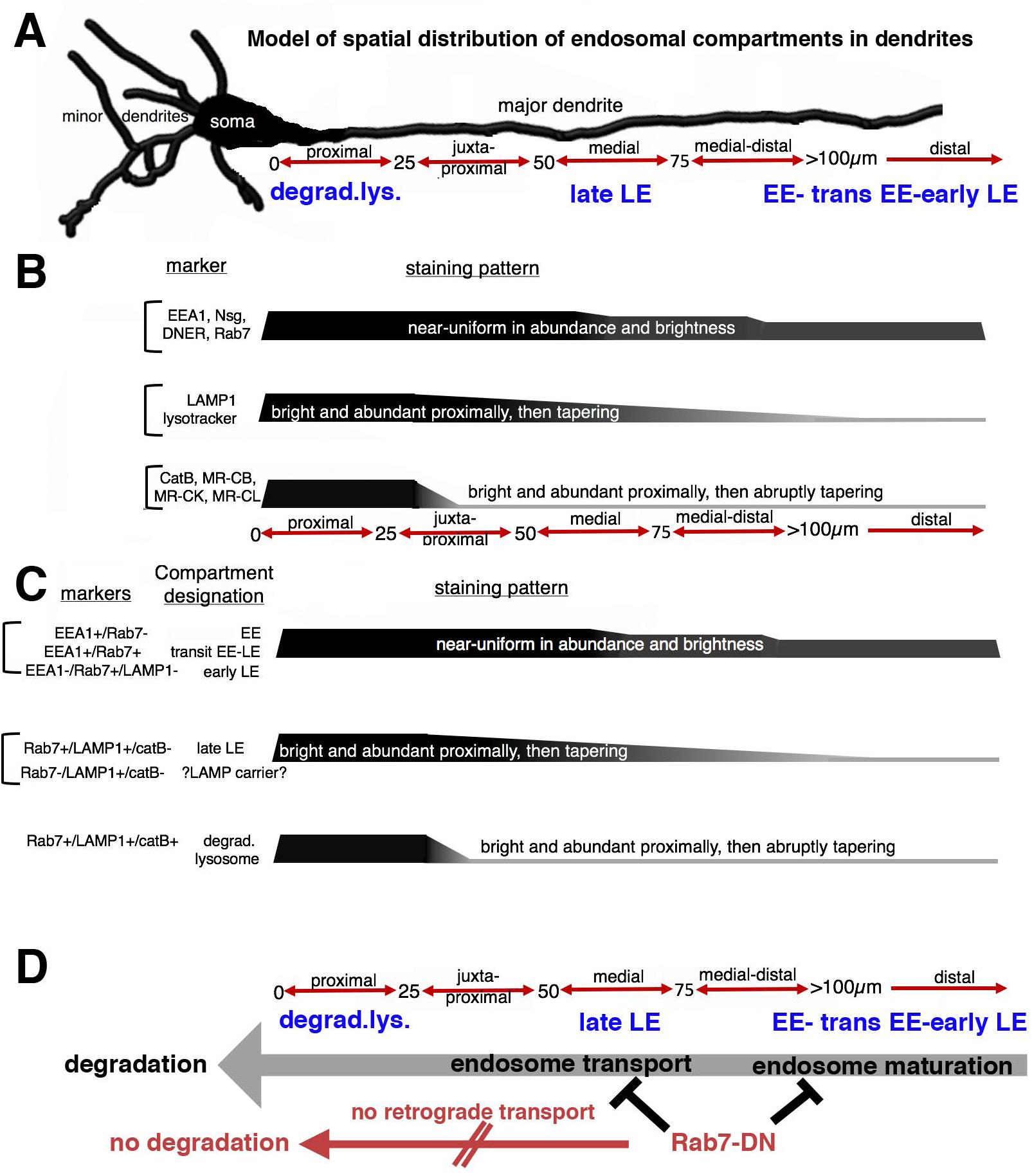
Summary of endosomal marker and compartment distribution in dendrites. (A) Diagram of major dendrite with the distance nomenclature (used throughout the paper) indicated along red arrows below. Compartment identities and their spatial appearance are indicated in blue. See (C) for markers used to define compartment identity. (B) Three distinct spatial distribution patterns can be observed for different markers, as indicated. (C) Model of spatial distribution patterns for distinct endosomal compartments. The compartment designations are indicated with the marker combinations used to define them indicated on the left. (D) Model for observed Rab7-DN effect on bulk degradation.

### Defining distinct endosomal compartments along dendrites: using multiple markers against endogenous proteins

The question of where lysosomes are located in neurons is surprisingly controversial. Several labs reported over many years that lysosomes (defined by LAMP1) tend to cluster in the neuronal soma (Cai et al., 2010; Maday and Holzbaur, 2016). Older work concluded that late endosomes (Rab7) as well as lysosomes (LAMP1 or endocytosed HRP) were clustered in the soma (Parton et al., 1992). Other works in contrast have described abundant lysosomes in axons and dendrites (Lee et al., 2011; Schwenk et al., 2014; Farias et al., 2017), using many of the same markers. Furthermore, Ehlers (2000) showed that AMPAR entered LAMP1-positive endosomes in dendrites and degraded rapidly after activity, but whether degradation took place locally was not clear. Along the same lines, recent work from several labs not only showed lysosomes in dendrites (Farias et al., 2017), but demonstrated activity-dependent transport of lysosomes to dendritic spines (Goo et al., 2017), or local secretion of lysosomal cathepsins at spines triggered by back-propagating action potentials (Padamsey et al., 2017).

These apparently contradictory findings are due in part to different definitions of what constitutes a lysosome as well as to the use of different markers to designate lysosomes. For instance, Rab7 and LAMPs are frequently interchangeably used to denote lysosomes (Schwenk et al., 2014). Similarly, late endosomes and lysosomes are often combined into a single category, and overexpressed markers (especially LAMP1-GFP) are used as stand-ins for revealing LE/lysosome localization (Farias et al., 2017). Depending on the questions asked, it can be useful to combine late endosomes and lysosomes into a single category. For our purposes of identifying where degradative capacity is localized in dendrites, we wanted to distinguish late endosomes from lysosomes. We thus define lysosomes as acidified LAMP1-positive compartments containing active cathepsins. We are employing the term “degradative lysosome” in this paper to designate these compartments. In order to definitively answer where different endosomal subpopulations reside in dendrites, we made use of a multitude of markers and determined the distribution of endogenous proteins, most often as combinations of three endogenous endosomal proteins simultaneously. In addition, we quantitatively determined how distribution of endosomal subpopulations changed with distance from the soma along dendrites. This has not been done quantitatively before and led to our discovery of a striking spatial gradient of endosomal populations along dendrites as well as to the identification of subpopulations of endosomes not previously described in dendrites, namely Rab7-positive/LAMP1-negative “early” late endosomes, and LAMP1-positive/Rab7-negative compartments of unknown function (Figure 9).

### A model for endosome maturation along dendrites

We find that early (EEA1-positive) endosomes and late (Rab7-positive) endosomes are widely distributed throughout soma and dendritic arbors. This agrees with our previous data that dendritic early endosomes largely do not move in dendrites and convert to late endosomes locally (Lasiecka et al., 2014; Yap et al., 2017). Late endosomes (Rab7-positive), on the other hand, have substantial motile populations in dendrites (Schwenk et al., 2014; Yap et al., 2017, and this work). We propose that it is the “early” late endosomes (Rab7-positive/LAMP1-negative) in medial and distal dendrites which are the major retrograde carrier for bulk degradative cargos back to the soma. Degradative lysosomes (CatB-positive/LAMP1-positive), in striking contrast, are overwhelmingly restricted to the soma and proximal portion of the major dendrite. In medial and distal portions of major dendrites, lysosomes are very sparse and contain comparatively lower levels of cathepsins (Figure 9). Degradative lysosomes are almost never found in the minor dendrites. Generally, the wider the dendrite, the farther from the soma can we detect degradative lysosomes along the major dendrite. We propose that Rab7 regulates retrograde transport of predegradative endosomes toward the soma/proximal dendrites as well as their maturation into degradation-competent LE/lysosomal compartments (Figure 9). Interestingly, unique LAMP-carriers have been identified in fibroblasts which deliver LAMP to late endosomes separately from cathepsins (Pols et al., 2013). Given the large extent of neuronal arbors and our discovery of steep spatial gradients of endosomal subpopulation in dendrites, where and how LAMPs and cathepsins are delivered to LEs/lys in dendrites is an open question. Furthermore, lysosome-related organelles containing low hydrolase activity are present in axons and accumulate in Alzheimer’s disease (Lee et al., 2011; Gowrishankar et al., 2015). Unlike our findings in dendrites though, Rab7 and LAMP1 are acquired more or less at the same time in distal axons (Lee et al., 2011). Similarly to our findings in dendrites, Rab7-positive/LAMP1-positive late endosomes in axons subsequently acquire CatB as they approach the soma (Lee et al., 2011).

### Rab7 is required for transport and bulk degradation of dendritic proteins

Our data support the conclusion that bulk degradation of dendritic cargos requires Rab7-dependent transport. Active Rab7 has a multitude of effectors which regulate many cellular processes, including recruitment of microtubule motors (dynein via RILP and kinesin-1 via FYCO1), recruitment of tether complexes (via HOPS complex subunits Vps39 and Vps41) for subsequent fusion, recruitment of retromer complex (via Vps35) for transport to the trans-Golgi network, and acidification (via RILP interactions with the V-ATPase) (Wang et al., 2011; Guerra and Bucci, 2016). Rab-DN is incapable of binding to any of its downstream effectors and its expression thus causes cessation of transport as well as disruption of LE/lys fusion and acidification in fibroblasts (De Luca et al., 2014). Consistent with findings in other cell types, overexpression of Rab7-DN in neurons resulted in a decrease in the movement of Nsg-endosomes and accumulation of Nsg/DNER-endosomes along dendrites after CHX treatment. These effects are presumably due to loss of RILP and HOPS recruitment to endosomes, but the detailed underlying molecular mechanisms will require additional study. We propose that bulk degradation of these short-lived dendritic membrane proteins occurs in somatic lysosomes after retrograde transport from dispersed Rab7-positive, pre-degradative “early” late endosomes in dendrites, rather than in distally localized, dendritic lysosomes (Figure 9). Since peripheral lysosomes in fibroblasts were found to be less acidified and less degradatively active than perinuclear ones (Johnson et al., 2016), our results suggest that there might be a similar spatial gradient in acidification and degradative capacity of LEs/lys in neurons. Whether these processes are similarly regulated in fibroblasts and neurons remains to be determined.

Interestingly, interference with Rab7 function leads to defects in dendrite maintenance in cultured neurons (Schwenk et al., 2014). Additionally, delayed dendrite growth was observed in the absence of the newly discovered Rab7 effector WDR91 (Liu et al., 2017), highlighting the physiological importance of Rab7 effector function and endosome maturation in dendrite maintenance. These dendritic phenotypes could be due to either disturbance of growth factor signaling from endosomes or to changes in bulk degradation or both (Casanova and Winckler, 2017). In addition, Rab7 function is required for long term depression in the cerebellum (Kim et al., 2017), likely by regulating degradation of glutamate receptors.

Rab7 also plays an important role in axons (Guerra and Bucci, 2016; Ponomareva et al., 2016). This is significant since mutations in Rab7 are genetically linked to Charcot-Marie Tooth disease 2B (CMT2B), a genetic disorder resulting in axonal neuropathy in the peripheral nervous system (Cogli et al., 2009). The axonal pathology might be due to altered NGF signaling from Rab7-containing TrkA signaling endosomes (Liu and Wu, 2017). Additional pathways are also regulated by Rab7 in axons (Cherry et al., 2013; Ponomareva et al., 2016), likely by affecting the half lives of additional cargos other than TrkA. Similarly, the final positioning of migratory neurons is regulated by Rab7 early in cortical development (Kawauchi et al., 2010), and N-cadherin is one of the proposed cargos regulated by Rab7 during neuronal migration.

### A specialized role for sparse degradative lysosomes in postsynaptic function?

We show using endogenous markers and rigorous quantification that there are only rare degradative lysosomes past the proximal dendrite (~ 25 *µ*m from the soma in cultured neurons). Recent work has shown the existence of biologically relevant LAMP1-positive compartments that might play a dedicated important role in synaptic function (Goo et al., 2017; Padamsey et al., 2017). Since fewer than 10% of spines contain LAMP1-positive compartments (Goo et al., 2017), we propose that neurons use somatic lysosomes for bulk degradation (this work), but might use a sparse set of synapse-associated LAMP1-compartments specifically dedicated to activity-dependent turnover of synaptic receptors (Goo et al., 2017). In addition, the sparse distal lysosomes might serve additional roles. Secretory lysosomes have been well described in other cell types (Luzio et al., 2014), but appear to also play important roles in neurons. For instance, Emptage and colleagues recently showed a role for dendritic secretion of CatB from lysosomes to promote activity-dependent spine growth (Padamsey et al., 2017).

In conclusion, we show that bulk degradation of dendritic membrane proteins overwhelmingly takes place in the soma or proximal dendrite. Since protein homeostasis needs to be maintained for neuronal health and is frequently at the center of neurodegenerative pathologies (Douglas and Dillin, 2010), it will be important to take the spatial constraints of bulk degradative potential into account.

## Materials and Methods

### Antibodies

Anti-L1CAM, clone 2C2, mouse monoclonal, 1:1000 WB, cat#ab24345, Abcam, RRID: <u>AB_448025</u>; anti-AMPAR2(GluR2), clone6C4, mouse monoclonal, 1:200 IF, 1:1000 WB, cat#MAB397, Millipore, RRID: <u>AB_11212990</u>; anti-GluR1, clone D4N9V, rabbit monoclonal, 1:200 IF, cat#13185, Cell Signaling; anti-syntaxin13, rabbit polyclonal, 1:1000 WB, cat#110 133, Synaptic System; anti-syntaxin7, sheep polyclonal, 1:1000 WB, cat#AF5478, R&D Systems, RRID: <u>AB_2239977</u>; anti-cathepsin B, goat polyclonal, 1:500 IF, 1:100 IHC, 1:1000 WB, cat#AF965, R&D Systems, RRID: <u>AB_2086949</u>; anti-cathepsin D, goat polyclonal, 1:1000 WB, cat#AF1029, R&D Systems, RRID: <u>AB_2087094</u>; anti-DNER, goat polyclonal, 1:500 IF, 1:1000 WB, cat#AF2254, R&D Systems, RRID: <u>AB_355202</u>; anti-LAMP2, rat monoclonal, 1:1000 WB, cat#ABL-93, DHSB, RRID: <u>AB_2134767</u>; anti-LAMP1, clone 1D4B, rat monoclonal, 1: 100 IHC, cat#1D4B, DHSB, RRID: <u>AB_2134500</u>. The information for antibodies against Nsg1, Nsg2, EEA1, LAMP1, Rab7, MAP2 and βIII tubulin can be found in Yap et al., 2017.

Secondary antibodies: As described in Yap et al., 2017, Alexa-dye coupled antibodies (Molecular Probes and Jackson ImmunoResearch) were used for Immunofluorescence. For Licor Odyssey Western blots, Jackson ImmunoResearch antibodies were used: Donkey anti mouse (680) #715-625-151, Donkey anti rabbit (790) #711-655-152, Donkey anti-sheep (680) #713-625-147, Donkey anti-goat (800) (Licor) #926-32214.

### Plasmids

*Nsg1-Emerald:* Addgene cat#54202, Michael Davidson lab; the mutation at residue 114 from D to G was corrected.

*Rab7GFP cat#12605, Rab7DN(T22N)-GFP cat#12660: Addgene, Richard Pagano lab mTagBFP2-Rab7:* Michael Davidson lab.

*GFP*: Clontech

mCherry-TFR-20 cat#55144: Addgene, Michael Davidson lab.

#### Neuronal cultures

Neuronal cultures were prepared from E18 rat hippocampi, as described in Yap et al 2017. All experiments were performed in accordance with institutional IACUC guidelines and regulations. Transfections were carried out with Lipofectamine 2000 (Invitrogen). Neurons were transfected with either GFP, GFP-Rab7 or GFP-Rab7-DN for 36-40 hours, and treated with/without cycloheximide (20μg/ml) for 4.5hrs. To investigate the effect of degradation inhibition on neuronal specific membrane proteins, neurons were treated with leupeptin (20μM) (Sigma-Aldrich) for 20hrs. All transfection experiments were repeated in five to ten independently derived cultures. Experiments using endogenous localization (Rab7, EEA1, cathepsinB, LAMP1) were repeated in at least three independent cultures.

#### Immunocytochemistry and endocytosis assay

All immunocytochemistry, endocytosis assay and image acquisition were done as described in Yap et al., 2017. For endocytosis assay in neurons treated with leupeptin, we treated DIV8 neurons with/without 20μM of leupeptin for 20 hrs, followed by endocytosis of anti-Nsg2, then chased the endocytosed antibody in the presence/absence of leupeptin for the indicated time. For quantification of colocalization and proteins abundance in dendrites, Z-stack images of fixed samples were acquired using Zeiss confocal 880.

#### Immunohistochemistry

Mice (Thy1-GFP M line) were maintained on a 12 h light/dark cycle and allowed *ad libitum* access to food and water. Experiments were conducted according to protocols approved by Drexel University Institute for Animal Care and Use Committee and Research Animal Resource Center.

Animals were given an intraperitoneal injection of ketamine/xylazine/acepromazine, and transcardially perfused with 20–30 ml of PBS followed by 60 ml of 4% paraformaldehyde. Brains were dissected and postfixed for 2–4 h, followed by cryoprotection in 30% sucrose, and stored at 4°C until ready for sectioning. Tissue was sectioned on a cryostat at 40 μm and serial sections were collected in 96 well plates containing 0.05% sodium azide, then stored at 4°C until ready for immunostaining. Tissue sections were processed as described in (Barford et al., 2017), except that the antigen retrieval step was omitted.

#### Functional labeling of lysosomes

Lysotracker Red (Lysotracker Red DND-99, 1:10000X, ThermoFisher, cat#L7528) was used to live label and track acidified compartments in transfected neurons using the manufacturer’s protocol. We used Magic Red cathepsin assay kits (dilution 1:1300, Immunochemistry Technologies, cat#937, #939 and #941) to live label active cathepsins B, K and L in transfected neurons according to the manufacturer’s protocol. DIV8 GFP-transfected neuronal cultures were incubated with 25μg/ml of DQ-BSA (DQ-BSA Red, ThermoFisher, Cat#D12051) and 25μg/ml of BSA-647 (BSA-Alexa 647 conjugate, ThermoFisher, cat#A34785) at 37C for 2hrs to live label protease-mediated hydrolysis of BSA together with endocytosed BSA-647 compartments. Images were captured live with the Axiocam 503 camera using Zen software (Zeiss) and processed identically in Adobe Photoshop.

#### Live Imaging and kymograph analysis

Live imaging was carried out as described in Yap et al., 2017. Briefly, neurons at DIV7 were transfected for 36 to 40 hrs with the following plasmids combinations: *Nsg1-mcherry* with *Rab7-GFP*, *Nsg1-mcherry* with *Rab7T22N-GFP, Nsg1-mcherry* with *GFP, mcherry-TFR* with *Rab7-GFP, mcherry-TFR* with *Rab7T22N-GFP,* and *mcherry-TFR* with *GFP.* Unconjugated transferrin (10μg/ml, ChromPure human transferrin, Jackson ImmunoResearch Laboratories, cat#009-000-050) was added to neurons before imaging. Kymographs were generated using KymographClear (Mangeol et al., 2016). Events were counted motile if vesicles movement was ≥ 2 μm and otherwise classified as stationary.

#### Western blot

Neurons at DIV10 were treated with leupeptin (20*µ*m) or cycloheximide (20 µg/ml) for the indicated time, washed several times with PBS, and harvested for subcellular fractionation using BioVision FractionPREP kit following the protocol described by the manufacturer. The cytosolic and membrane fractionated samples were subjected to SDS-PAGE and western blot analysis.

#### Image analysis and quantification

##### Somatic intensity analysis

Ten fields for each condition were analyzed the Imaris 8.4 image analysis package. In each field the soma of discernable cells (six to ten per field) were outlined by hand to exclude the nucleus. For the image as a whole the average intensity of each marker in these outlined regions was calculated. The values for each field were normalized to the mean of the respective marker minus leupeptin. Statistics were calculated from these average values of the ten images. Using Imaris 9.0, for each field of view principle dendrites were identified that could be traced cleanly without interference from other dendrites (total of 17 dendrites). They were then segmented in 25 *µ*m sections stating at the edge of the cell body with the first labeled proximal, the middle section medial, and the last section distal. For each marker a threshold level was determined and used across all images. Surface objects were created using the established threshold.

##### Abundance and volume analysis

For the abundance analysis the number of objects obtained were counted and plotted. Each dot in the resulting graph represents the number of objects in the specified region of one dendrite. For the volume analysis the average volume of all objects in a segment were plotted with each dot in the resulting graph representing the specified region in one dendrite.

##### Quantification of Nsg2 endocytosis

Images were quantified for fluorescence intensity in the soma and in the dendrite at time zero and following a 4h chase using FIJI. Background levels were subtracted for each measurement. For the dendrites, intensity of a line (line width 15) an average length of 22-30 *µ*m was drawn at least 15 *µ*m from the soma. The intensity along that line was measured and then the background subtracted from it. Values were normalized and the Kruskal-Wallis test was used to determine significance between groups.

## Summary of Supplementary Figures

Suppl. Fig. 1 shows Rab7-GFP in combination with endogenous EEA1 and LAMP1.

Suppl. Figure 2 shows co-imaging of endosomes with LysotrackerRed and MRCatB.

Suppl. Figure 3 shows imaging of active cathepsins K and L (MR-CatK and MR-CatL).

Suppl. Figure 4 shows Rab7-DN inhibits degradation of DNER.

## Acknowledgments

We thank all members of the Winckler Laboratory for constructive engagement throughout the duration of this work and critical reading of the manuscript. We gratefully acknowledge the extremely valuable input from Drs. David Castle, Jim Casanova, and Ryan D’Souza through many discussions and reading of drafts. This work was supported by NIH grant R01NS083378 (to BW).

